# Dual regulation of SLC25A39 by AFG3L2 and iron controls mitochondrial glutathione homeostasis

**DOI:** 10.1101/2023.11.06.565855

**Authors:** Xiaojian Shi, Marisa DeCiucis, Kariona A. Grabinska, Jean Kanyo, Adam Liu, Tukiet Lam, Hongying Shen

**Affiliations:** Cellular and Molecular Physiology Department, Yale School of Medicine, New Haven, CT, USA; Systems Biology Institute, Yale West Campus, West Haven, CT, USA; Keck MS & Proteomics Resource; Yale/NIDA Neuroproteomics Center, Yale University, New Haven, CT, USA; Amity High School, Woodbridge, CT, USA

## Abstract

Organelle transporters define metabolic compartmentalization and how this metabolite transport process can be modulated is poorly explored. Here, we discovered that SLC25A39, a mitochondrial transporter critical for mitochondrial glutathione uptake, is a short-lived protein under dual regulation at the protein level. Co-immunoprecipitation mass spectrometry and CRISPR KO in cells identified that mitochondrial *m*-AAA protease AFG3L2 is responsible for degrading SLC25A39 through the matrix loop 1. SLC25A39 senses mitochondrial iron-sulfur cluster using four matrix cysteine residues and inhibits its degradation. SLC25A39 protein regulation is robust in developing and mature neurons. This dual transporter regulation, by protein quality control and metabolic sensing, allows modulating mitochondrial glutathione level in response to iron homeostasis, opening new avenues to explore regulation of metabolic compartmentalization. Neuronal SLC25A39 regulation connects mitochondrial protein quality control, glutathione and iron homeostasis, which were previously unrelated biochemical features in neurodegeneration.

## Introduction

Mitochondrial metabolite transporters^1-3^ define metabolic compartmentalization between mitochondria and rest of the cell^4^, and transport dysregulation is associated with many pathological conditions such as neurodegenerations, metabolic syndromes and cancers. Despite the recent discoveries of protein identities that facilitate metabolite transport, the mechanism by which transporter proteins can be regulated remain unexplored.

Given the critical role of mitochondrial redox metabolism and oxidative stress^5-8^ in physiology, we focus on the regulation of mitochondrial transport for glutathione, the most abundant endogenous antioxidant. Glutathione^9^ critically supports redox signaling and biosynthesis of iron-sulfur (Fe-S) cluster cofactors inside the mitochondrial matrix^10-12^. Because glutathione is exclusively synthesized in the cytosol, how glutathione enters the mitochondrial matrix remained a decades-old mystery until the recent identification that a previously uncharacterized mitochondrial metabolite transporter, SLC25A39 (A39), is critical for mitochondrial glutathione transport^13-15^. A39 belongs to the <u>S</u>o<u>L</u>ute <u>C</u>arrier 25 (SLC25) family, the largest transporter family that is responsible for translocating diverse metabolite ligands across the mitochondrial inner membrane^1-3^. Despite the relatively small mitochondrial volume, mitochondrial glutathione accounts for 10-15% of the total cellular glutathione pool^10^. We therefore hypothesize that a direct regulation of A39 could have a major impact on glutathione compartmentalization and cellular metabolism.

## Results

### SLC25A39 protein regulation depends on GSH and the matrix cysteine residues

To explore the regulatory mechanism of A39, we performed amino acid sequence analysis^14,16^ and Alphafold structure prediction^17^ of A39. In addition to the typical structural fold, the threefold pseudo-symmetrical, six transmembrane α-helices surrounding the central transport cavity, the analysis revealed that A39 contains an extra mitochondrial matrix loop between transmembrane domain 1 and 2 with a low structural prediction confidence score (Fig. 1A). This loop 1 of amino acids 41-105 contains four matrix side surface-exposed, thiol Cysteine (Cys) residues^18,19^ (Fig. 1A), which exceeds Cys frequency in proteins at around 1.7%^20^. C-terminal FLAG-tagged A39 construct with all four cysteines mutated to alanine, A39^4CA^-FLAG, can fully restore mitochondrial GSH depletion in the A39 CRISPR KO cells similar to the wild type A39-FLAG (Fig. 1C), suggesting that the A39^4CA^ mutant is an active transporter, while another cysteine mutant of the substrate binding site A39^C334S^-FLAG can only rescue partially (Fig. 1C). However, A39^4CA^ -FLAG completely abolished A39 protein upregulation upon pharmacological inhibition of glutathione biosynthesis using buthionine sulphoximine (BSO) (Fig. 1B), a regulation specific to A39 protein at the post-transcriptional level (Fig. S1A-C) that was previously reported with an unknown mechanism^13^.

**Figure 1.**
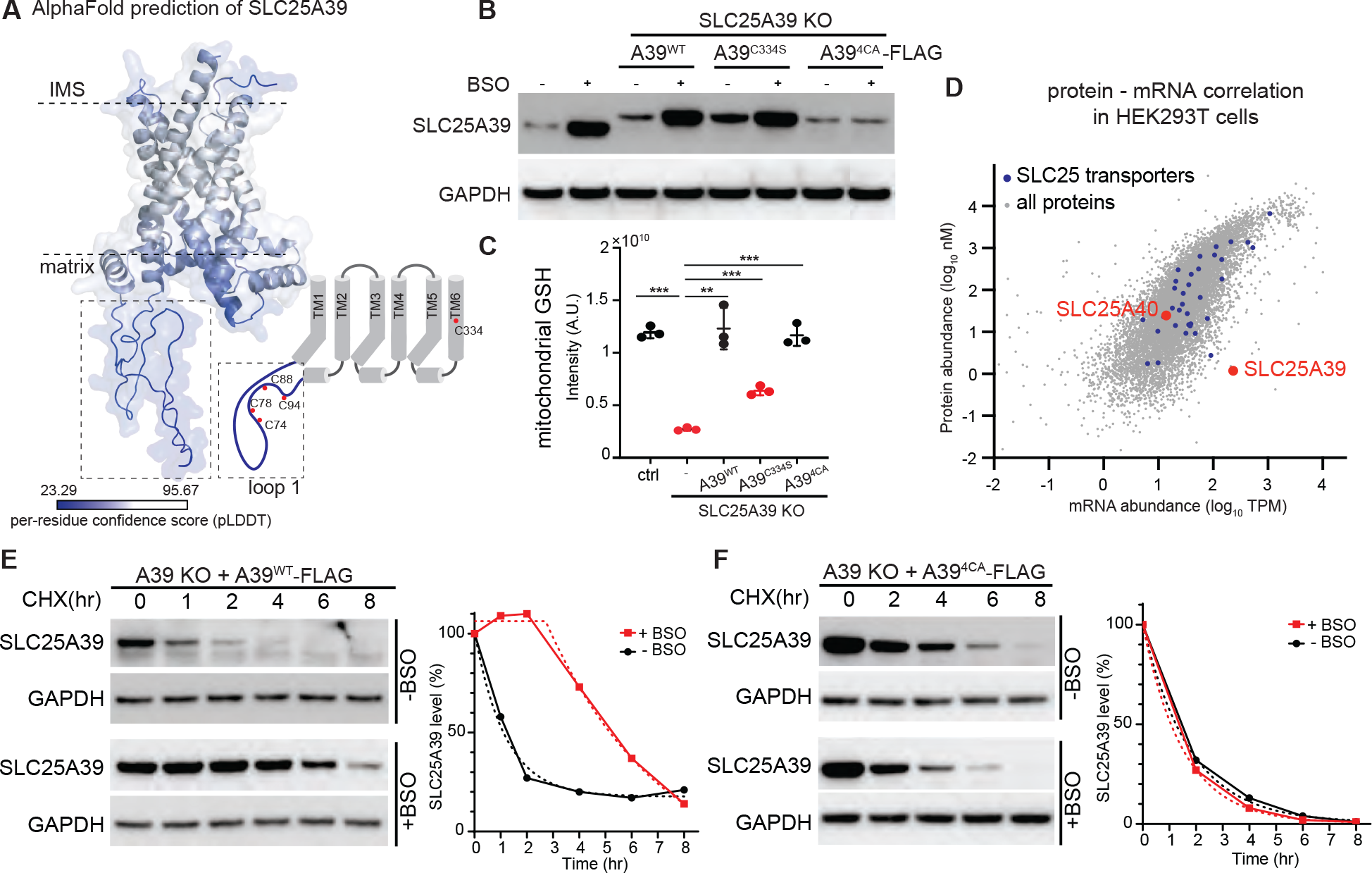
SLC25A39 (A39) is a short-lived protein, and the degradation is regulated by GSH and the matrix cysteine residues. (A) AlphaFold prediction of A39 structure revealed a unique cysteine-rich matrix loop 1. (B) Western blot of endogenous A39 and ectopically expressed FLAG-tagged A39^WT^, substrate binding pocket mutant A39^C334S^, and matrix loop 1 four cysteine mutant A39^4CA^ in the K562 cells upon GSH depletion by BSO treatment (1mM, 2d). GAPDH, loading control. (C) Mitochondrial GSH level assayed by HA-MITO tag immunoisolation followed by MS-based metabolite profiling in the control and clonal A39 CRIPSR KO K562 cells expressing the indicated A39 constructs. Statistical significance was calculated using two-tailed *t* test. Significance level was indicated as ^***^ *p* < 0.001, ^**^ *p* < 0.01. Data are expressed as mean ± SD. (D) Scatter plot showing the correlation of protein abundance (log_10_nM) and mRNA abundance (Log_10_ TPM) for all coding genes expressed in the HEK293 cells (OpenCell, CZI). Blue dots, SLC25 transporter proteins. A39 and A40 are highlighted in red. (E-F) Western blot showing A39-FLAG (E) and A39^4CA^-FLAG (F) protein level in the clonal A39 CRISPR KO K562 cells treated with CHX (cycloheximide, 150 μg/ml) for the indicated times, either under basal condition (-BSO) or with BSO (1mM and with 2 d prior treatment). GAPDH, loading control. A39 band intensity was quantified and modeled using the non-linear fitting with one phase decay (A39, -BSO; A39^4CA^, -BSO and +BSO) and plateau followed by one phase decay (A39, +BSO).

### SLC25A39 is a short-lived protein and degradation is regulated by cysteine residues

Because ectopically expressed recombinant A39-FLAG protein can only be expressed as a level similar to that of the endogenous A39 (Fig. 1B), we suspected that A39 protein might be tightly maintained at a low level under basal condition that is then alleviated upon GSH depletion by BSO. Indeed, we first identified that human A39 protein level is extremely low in comparison to the A39 mRNA level, which stands out as a proteome-wide outlier of protein-mRNA correlation in the HEK293T cells (CZI Open Cell^21^, Fig. 1D) — the majority of the mitochondrial SLC25 transporters exhibit correlated protein and mRNA levels (Fig. 1D). Because all inner membrane SLC25 transporters are imported through a common TIM22 import mechanism, we suspected that A39 protein stability is regulated. We then performed cycloheximide (CHX) chase experiments and discovered that pre-synthesized A39 protein indeed exhibited a very short half-life < 2 hrs under basal condition (Fig. 1E). A39^4CA^ protein exhibited a similar short half-life (Fig. 1F). This half-life is drastically shorter than the majority of mitochondrial proteins^22,23^ of approximately median half-life at 87 h^24^. BSO significantly extended A39’s half-life (Fig. 1E), but failed to extend the short half-life of A39^4CA^ (Fig. 1F). This suggests that under basal condition, both pre-synthesized wild-type A39 and A39^4CA^ mutant were quickly degraded, and BSO treatment inhibited A39 degradation through a mechanism that depends on the matrix cysteines. These two regulatory mechanisms of A39, both fast degradation and BSO-mediated stabilization, are yet to be explored.

### Mitochondrial *m*-AAA protease AFG3L2 is responsible for A39 degradation

To characterize the quality control pathway that degrades A39, we combined co-immunoprecipitation mass spectrometry (MS) and CRISPR KO in cells to identify its regulatory machinery. We first selected K562 clonal cells ectopically expressing relatively high level of A39-FLAG and a mitochondrial tag (HA-MITO, 3xHA-eGFP-OMP25^25,26^) from the clonal A39 CRISPR KO cells (Fig. 2A and S2A). We treated the cells with BSO for 2 days to increase the A39-FLAG protein level (Fig. S2A), rapidly immuno-isolated the mitochondria via HA-MITO using anti-HA magnetic beads, and then immunoprecipitated A39 binding partners using anti-FLAG resin in a mild 1% digitonin-containing lysates (Fig. 2A). Wild type K562 cell expressing HA-MITO tag was used as the control. Silver staining of the immunoprecipitants revealed an enrichment of A39-FLAG protein that migrated around 40 kD, confirmed by MS (Fig. 2A). One major band around 60kD that is specific to A39-FLAG IP was identified as HSPD1/HSP60 by MS (Fig. 2A), an abundant mitochondrial matrix chaperonin that facilitates folding and refolding of matrix proteins^27^. HSPD1 enrichment suggested that the matrix portion of A39 protein may be flexible and misfolded, however, because HSPD1 CRISPR KO did not affect A39 protein level under basal condition (Fig. 2C), we did not follow up here. We then performed MS analysis of the entire gel lane from the A39-FLAG immunoprecipitants and control to identify specific A39 binding partners (Supplementary Data 1). We categorized the hits into pathways relevant to mitochondrial quality control and A39-related metabolite sensing mechanisms that include mitochondrial proteases, matrix chaperones, mitophagy-related, and iron-related proteins (Fig. 2B). We then performed CRISPR to knock out 11 representative protein hits from these pathways to investigate the impact on A39 protein level, both under basal condition and BSO treatment (Fig. 2C). Strikingly, CRISPR KO of only two proteins dramatically stabilized A39 under basal condition, AFG3L2 and ABCB7 (Fig. 2C, red), among which AFG3L2 KO upregulated basal A39 protein level to a much higher level and A39 level in the AFG3L2 KO was no longer sensitive to BSO treatment (Fig. 2C).

**Figure 2.**
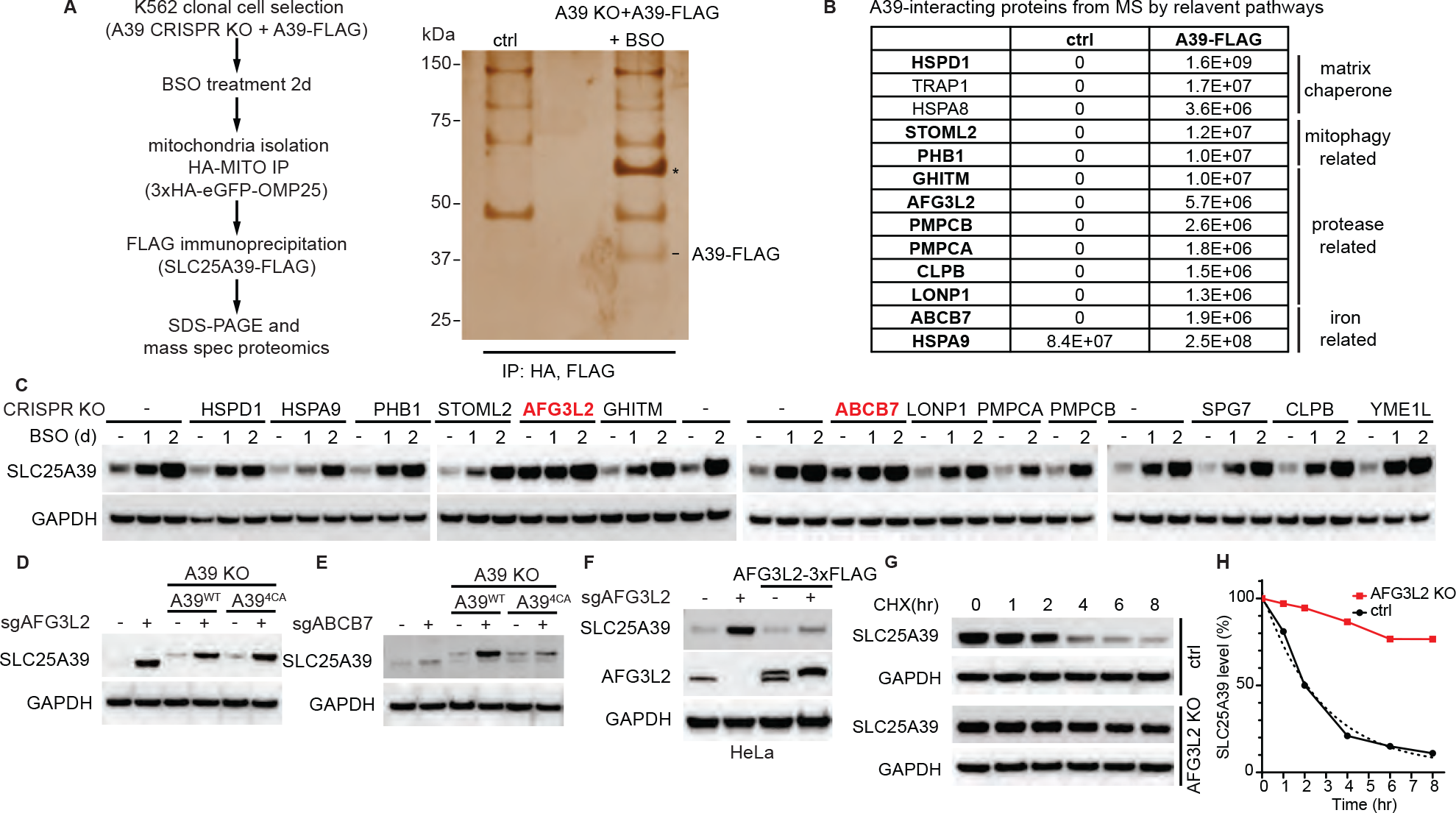
Co-immunoprecipitation mass spectrometry and CRISPR KO identify the mitochondrial *m*-AAA protease AFG3L2 in A39 degradation. (A) Left, flow chart showing the experimental procedure to identify putative A39-interacting regulatory proteins. Right, silver stained SDS-PAGE of anti-FLAG immunoprecipitants from the control and A39-FLAG mitochondrial lysates. Wild type K562 cells expressing 3HA-eGFP-OMP25 were used as control. A39-FLAG protein identified by MS is labeled; *, a specific binding partner HSPD1 identified by MS. (B) Categorization of specific A39-interacting protein hits into relevant quality control pathways. Protein level intensity based on top three peptides was shown in the control and A39-FLAG co-immunoprecipitants. Bold, hits that were followed up by CRISPR KO. (C)Western blot of A39 from CRISPR KO K562 cells using sgRNAs targeting indicated A39-interacting proteins, and related mitochondrial proteases. Cells were analyzed under basal condition and BSO (200 μM) treatment for either 1 d or 2 d. Two CRISPR KO lines that increased basal A39 level were highlighted in red. GAPDH, loading control. (D-E) Western blot of endogenous A39, ectopically expressed FLAG-tagged A39^WT^ and A39^4CA^ in the AFG3L2 CRISPR KO cells (+ sgAFG3L2, D) and ABCB7 CRISPR KO cells (+sgABCB7, E). (F) Western blot of endogenous A39 in HeLa cells upon AFG3L2 CRISPR KO and re-expressing AFG3L2-3xFLAG. GAPDH, loading control. (G) Western blot of the endogenous A39 protein in the K562 cells treated with CHX (cycloheximide, 150 μg/ml) for the indicated times, with or without AFG3L2 CRISPR KO. GAPDH, loading control. (H) A39 band intensity was quantified and modeled by the non-linear fitting with one phase decay.

AFG3L2 is the subunit of the mitochondrial *m-*AAA proteases in the inner membrane with catalytic sites facing the matrix^28^, and its mutations cause neurological disorders including dominant spinocerebellar ataxia (SCA28)^29^ and Spastic ataxia 5, autosomal recessive (SPAX5)^30^. AFG3L2 CRISPR KO increased the basal level of endogenous A39, A39-FLAG and A39^4CA^-FLAG (Fig. 2D), suggesting its role in the quality control of A39 (and A39^4CA^) under basal condition. ABCB7 is a mitochondrial inner membrane protein and a putative transporter for certain iron-sulfur cluster (Fe-S) species^31^. Because ABCB7 CRISPR KO only increased wild-type A39-FLAG but not A39^4CA^-FLAG (Fig. 2E), we suspected that ABCB7 might function in the second branch of the regulation, the BSO-mediated A39 stabilization that depends on the matrix cysteines.

The quality control regulation of A39 protein is specific to AFG3L2. Two hexametric *m-*AAA proteases exist in human, homo-oligomeric AFG3L2 complexes and hetero-oligomeric complexes of the homologous AFG3L2 and SPG7^32,33^. Together with *i-*AAA protease active in the intermembrane space YME1L and other membrane-bound peptidases, they play a central role for inner membrane protein quality control. Notably, our co-immunoprecipitation MS did not recover SPG7 and YME1L, and CRISPR KO of SPG7 and YME1L did not affect A39 protein level (Fig. 2C), suggesting a specificity by AFG3L2.

CHX chase experiment identified that AFG3L2 CRISPR KO completely abolished A39 degradation beyond 8 hrs (Fig. 2G and H) --a complete stabilization upon AFG3L2 loss is different from the moderate BSO-mediated stabilization (Fig. 1E). This suggests that A39 is a specific, short-lived substrate of AFG3L2, and other mitochondrial proteases cannot compensate and substitute for A39 proteolytic regulation. We validated these findings in HeLa cells, in which AFG3L2 CRISPR KO cells also upregulated A39 level that can be reduced by re-expressing AFG3L2-3xFLAG (Fig. 2F). We therefore concluded that AFG3L2 is necessary for degrading A39 under basal condition and this function cannot be substituted by other mitochondrial proteases.

### A39 degradation is targeted through the matrix loop 1, which fine-tunes mitochondrial glutathione level

We then tested whether the matrix loop 1 of A39 is responsible for A39 degradation by AFG3L2. We generated two A39 fusion constructs by swapping the long A39 loop 1 to the corresponding loop of either the homologous transporter SLC25A40 (A40), A39^A40L1^ -FLAG, or to that of the dicarboxylate SLC25 transporter SLC25A11 (A11), A39^A11L1^-FLAG (Fig 3A and B). As expected, due to the shorter sequences, A39^A11L1^-FLAG and A39^A40L1^-FLAG migrated at lower molecular weights than A39-FLAG (Fig. 3C). Importantly, A39^A11L1^-FLAG and A39^A40L1^ -FLAG were expressed at a much higher level than A39-FLAG under basal condition (Fig. 3C), and AFG3L2 CRISPSR KO cannot further stabilize these two fusion proteins (Fig. 3C), suggesting they were resistant to degradation. A lack of AFG3L2-mediated degradation for the loop 1-swapping A39 fusion constructs strongly suggested that A39’s matrix loop 1 is responsible for AFG3L2 recognition and degradation.

**Figure 3.**
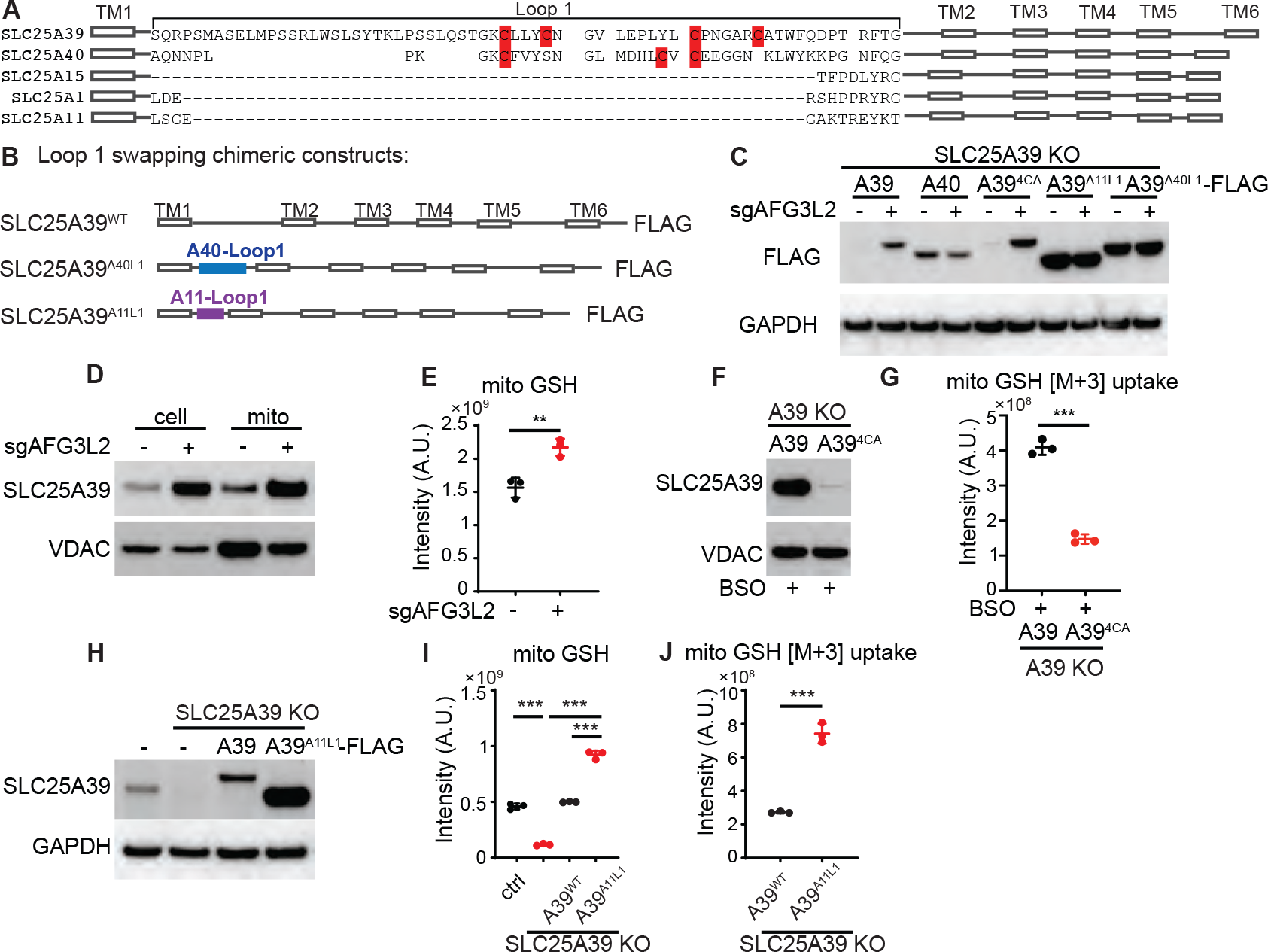
A39 degradation is targeted through the matrix loop 1, which controls mitochondrial glutathione uptake and steady state level. (A) Sequence alignment showing matrix loop 1 of A39, SLC25A40, and other representative SLC25 transporters SLC25A15, SLC25A1 and SLC25A11. (B)Diagram showing Loop 1 swapping chimeric constructs. (C) FLAG western blot for the indicated constructs in the clonal A39 CRISPR KO K562 cells, upon AFG3L2 CRISPR KO (+sgAFG3L2). GAPDH, loading control. (D) Western blot of endogenous A39 in the whole cell lysates and mitochondrial lysates from the control and AFG3L2 CRISPR KO K562 cells. Mitochondrial protein VDAC, loading control. (E) Mitochondrial GSH level assayed by HA-MITO tag immunoisolation followed by MS-based metabolite profiling in the control and AFG3L2 CRIPSR KO K562 cells, normalized by total protein abundance. (F) Western blot of A39 from BSO (1mM, 2 d)-treated clonal A39 CRISPR KO cells expressing either wild type A39 and A39^4CA^-FLAG. VDAC, loading control. (G) GSH uptake (1mM labeled GSH + 4mM unlabeled GSH, 15 min at room temperature) into HA-MITO immunoisolated mitochondria from BSO-treated clonal A39 CRISPR KO cells expressing either wild type A39 or A39^4CA^-FLAG. (H) Western blot of A39^WT^ and A39^A11L1^-FLAG in the clonal A39 CRISPR KO cells. (I) Mitochondrial GSH level assayed by HA-MITO tag immunoisolation followed by MS-based metabolite profiling in control, A39 CRISPR KO, and A39 CRISPR KO re-expressing A39^WT^, or A39^A11L1^-FLAG, normalized by NAD^+^ level. (J) GSH uptake (5mM labeled GSH, 15 min at room temperature) into isolated mitochondria from A39 CRISPR KO cells expressing A39^WT^ and A39^A11L1^-FLAG. All statistical significance was calculated using two-tailed *t* test. Significance level was indicated as ^***^ *p* < 0.001, ^**^ *p* < 0.01. Data are expressed as mean ± SD.

We then explored the functional consequence of A39 upregulation on mitochondrial glutathione level using three different experimental conditions. First, we performed mitochondrial metabolite profiling in the AFG3L2 CRISPR KO cells upon rapid mitochondrial immunoisolation using HA-MITO tag and discovered that the KO mitochondria lacking the protease had a higher A39 protein level as expected (Fig. 3D), and increased GSH level (Fig. 3E). Second, we immunoisolated mitochondria from BSO-treated A39 CRISPR KO cells expressing either A39-FLAG or A39^4CA^-FLAG where only A39-FLAG can be upregulated but not A39^4CA^-FLAG (Fig. 3F); and then performed *in vitro* mitochondria-based uptake assay using stable isotope labeled GSH [M+3] (GSH-[glycine-^13^C2, ^15^N]) for 15 min. The uptake assay revealed significantly higher GSH uptake in the wild type A39-FLAG expressing cells (Fig. 3G), despite A39^4CA^-FLAG is active (Fig. 1B). Third and specifically, we compared mitochondrial glutathione level in the A39 KO cells overexpressing A39-FLAG or A39^A11L1^-FLAG, in which A39^A11L1^-FLAG is expressed at a much higher level than A39-FLAG (Fig. 3H). Indeed, while A39-FLAG can rescue mitochondrial glutathione depletion in the A39 KO cells to the control level, A39^A11L1^-FLAG further increased mitochondrial GSH higher than that of the control level (Fig. 3I). Labeled GSH mitochondrial uptake assay from these cells also confirmed a high mitochondrial glutathione uptake rate (Fig. 3J). Therefore, we concluded that a regulated A39 transporter protein level directly controls mitochondrial matrix glutathione level.

### SLC25A39 degradation is inhibited by a coordinated sensing of mitochondrial iron-sulfur cluster by the matrix cysteines

What is the mechanism that stabilizes A39 protein during BSO treatment? We first tested and ruled out the possibility that BSO affects AFG3L2 activity, because BSO treatment does not affect level of NDUFA9 (Fig. 4A), whose degradation depends on AFG3L2 in the mammalian cells^34^. Because previous studies of A39 identified a coupling between A39 and mitochondrial iron homeostasis^13,14,35,36^, we hypothesized a BSO-mediated, A39’s matrix-cysteine-dependent metabolic sensing mechanism. Consistent with this notion, our CRISPR KO assay (Fig. 2C) already observed that CRISPR KO of the putative mitochondrial Fe-S cluster transporter ABCB7 stabilized A39 protein in a similar manner that depends on A39’s matrix cysteines (Fig. 2E). Following up on this, we deployed three strategies to perturb iron homeostasis to investigate the impact on A39 protein level (Fig. 4B). First, depleting cellular iron using an iron chelator deferoxamine (DFO), which is sensed by increased cytosolic iron-sensing IRP2 level (Fig. S3A and S3B), diminished BSO-mediated A39 protein upregulation (Fig. 4C and S3B). Second, depleting mitochondrial iron by double CRISPR KO of the two mitochondrial iron transporters, SLC25A28 and SLC25A37 (Fig. S3C and S3D), completely abolished BSO-mediated A39 protein upregulation (Fig. 4D). Third, inhibiting Fe-S cluster biosynthesis chaperone by CRISPR KO of HSCB or GLRX5 (Fig. S3E and S3F) significantly dampened BSO-mediated A39 protein upregulation (Fig. 4E). Then, to investigate the contribution of individual matrix cysteines in A39, we generated four FLAG-tagged single cysteine mutants, A39^C74A^, A39^C78A^, A38^C88A^ and A39^C94A^. While all four mutants can be expressed similar to the wild-type level under basal condition, all four mutants exhibited reduced stabilization upon BSO treatment (Fig. S3G), suggesting a coordinated regulation through all four cysteines.

**Figure 4.**
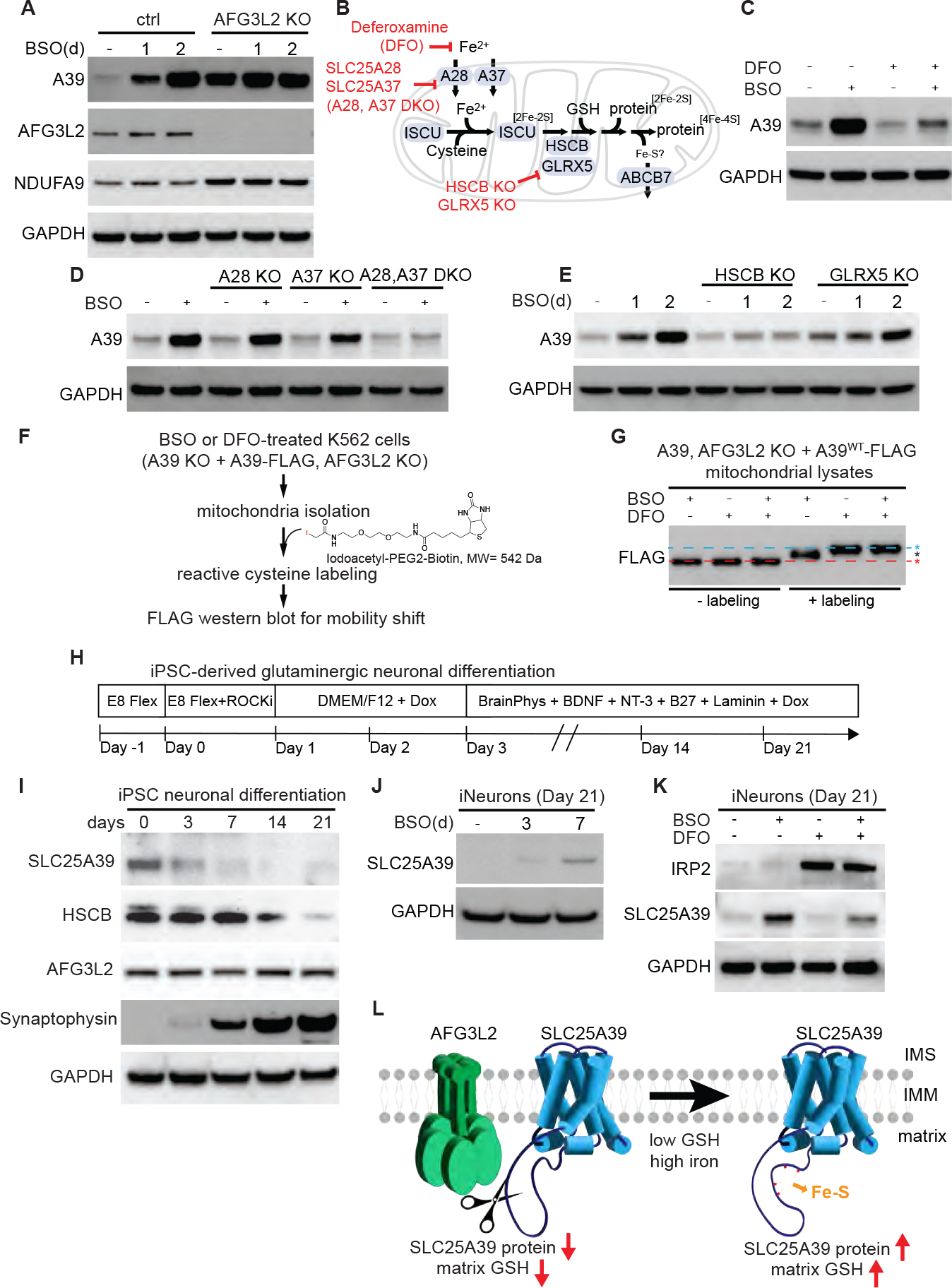
A39 is stabilized through a coordinated sensing of mitochondrial iron-sulfur cluster by the matrix cysteines. A39 regulation occurs in neurons. (A) Western blot of A39 showing BSO treatment (1mM, 2d) increased A39 in the control K562 cells but did not affect NDUFA9 level, while the basal level of A39 and NDUFA9 were increased in the AFG3L2 CRISPR KO K562 cells. (B) Diagram of iron manipulation strategies. (C)Western blot of A39 in K562 cells treated with either or a combination of BSO (1mM, 2d) and DFO (20 μM, 2d). (D) Western blot of A39 with and without BSO (1mM, 2d) in the CRISPR KO K562 cells of either or both mitochondrial iron transporters SLC25A28 and SLC25A37. (E) Western blot of A39 upon BSO (1mM) treatment for 1 and 2d in the CRISPR KO K562 cells of HSCB or GLRX5. (F) Diagram of A39 cysteine labeling and mobility shift assay. Cells were treated with either or a combination of BSO (100 μM, 2d) and DFO (deferoxamine, 20 μM, 2d). (G) Anti-FLAG western blot and mobility shift assay for A39-FLAG in the indicated conditions, without or with cysteine labeling. (H) Diagram of neuronal differentiation from the iPSC line, KOLF2.1J cells expressing dox-inducible *NGN2* in the safe locus. (I) Western blot of A39, HSCB, AFG3L2, and Synaptophysin, a synaptic marker from the indicated days (H) during neuronal differentiation. GAPDH, loading control. (J) Western blot of A39 in the mature iNeurons (differentiation Day 21) upon 1 mM BSO treatment for 3 and 7 d. GAPDH, loading control. (K) Western blot of A39 and IRP2 in the mature iNeurons (differentiation Day 21), treated with either or a combination of BSO (1mM, 7d) and DFO (20 μM, 7d). GAPDH, loading control. (L) A39 regulation model.

To assay whether the four matrix cysteines might be directly involved in iron sensing, we applied reactive cysteine labeling reagent, iodoacetyl-PEG2-biotin (MW = 542 Da), to label cysteine residues of A39^WT^-FLAG in the A39 and AFG3L2 KO K562 cell mitochondrial lysates, and assayed iron-dependent labeling efficiency through A39-FLAG mobility shift (Fig. 4F). Because A39 protein contains three additional cysteine residues besides the four matrix cysteines, we observed a modest mobility shift of the labeled A39-FLAG from the BSO-treated condition, corresponding to a partial labeling (Fig. 4G, black asterisk). Importantly, cellular iron depletion by either DFO or BSO + DFO treatment allowed A39-FLAG to be maximally labeled and to migrate at a higher molecular weight (Fig. 4G, cyan asterisk), suggesting more cysteine residues of A39 became exposed upon iron depletion and reactive to labeling. The mobility shift reflects cysteine labeling, because A39-FLAG from mitochondrial lysates prior to labeling migrated at the similar molecular weight (Fig. 4G, red asterisk) and the protein abundance is comparable due to AFG3L2 KO. We therefore concluded that A39’s four matrix cysteines are responsible for coordinated sensing of matrix iron homeostasis, protecting A39 from degradation.

### A39 regulation in the neuronal cells

Because glutathione, iron, and mitochondrial dysregulation have all been implicated in neurological disorders, we then explored if this A39 protein regulation mechanism also occurs in the neuronal cells. We chose the human induced pluripotent stem cell (iPSC) KOLF2.1J cell-derived glutamatergic neurons utilizing Tet-On doxycycline (Dox)-inducible NGN2 expression (Fig. 4H)^37-39^. The neuronal differentiation was robust (>95%) based on neuronal morphology and they were considered mature iNeurons on differentiation Day 21 (Fig. S3H). Western blotting for synaptic marker Synaptophysin confirmed synaptic development (Fig. 4I). We observed a time-dependent reduction of A39 protein level during neuronal differentiation (Fig. 4I), which is accompanied by a decline of the Fe-S cluster machinery HSCB (Fig. 4I), suggesting a coordinated regulation of mitochondrial glutathione and Fe-S cluster during differentiation. The AFG3L2 protein level remained unchanged (Fig. 4I). A39 reduction in neurons is likely due to reduced iron sensing that activates degradation, because BSO treatment in the mature iNeurons dramatically upregulated A39 protein level (Fig. 4J and 4K), which can be dampened by DFO treatment (Fig. 4K). The identification of A39 regulation in the neuronal cells might suggest a regulated neuronal glutathione compartmentalization in the context of brain disorders.

## Discussion

Here, we reported a brand-new dual regulatory mechanism acting on a mitochondrial transporter protein, by both protein quality control and metabolic sensing, which directly controls mitochondrial glutathione level in response to iron homeostasis (Fig. 4L). The proteolytic regulation of A39 is specific to the mitochondrial *m*-AAA protease AFG3L2, and cannot be substituted by other mitochondrial proteases that also act around the inner mitochondrial membrane^40,41^. A direct sensing of mitochondrial iron homeostasis inhibits A39 degradation, further enabling a specific transporter regulation and coordinated metabolism of two essential and reactive redox metabolites, glutathione and iron, a finding independently reported by Liu et al.^42^. We further went on to reveal this A39 regulation in the developing and mature neurons. This unexpected link between mitochondrial glutathione and *m*-AAA protease AFG3L2 might lead to new pathophysiological mechanisms under a diverse clinical spectrum of AFG3L2 mutation-associated neurodegenerative disorders that range from ataxia to parkinsonism^29,43^. This dual regulatory mechanism of SLC25A39 transporter, by AFG3L2 and iron, further advances our understandings of metabolic regulation through protein quality control and metabolite sensing, opening new avenues to explore the regulation of metabolic compartmentalization in cellular metabolism.

---

**Figure S1.**
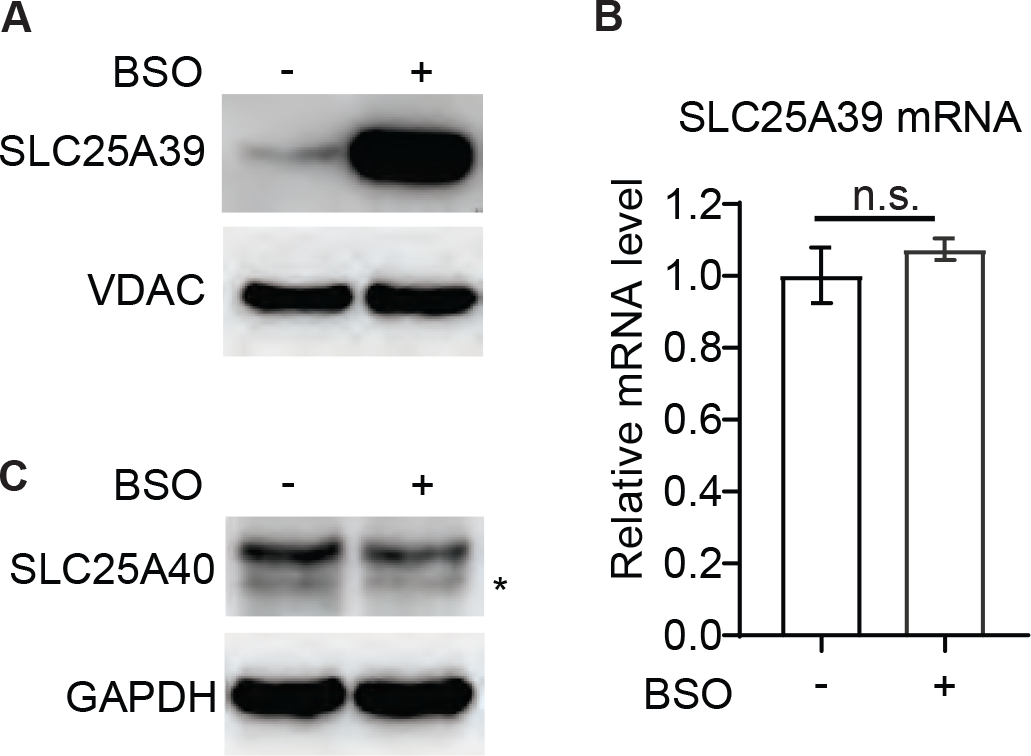
Western blot of A39 (A) and the paralogous SLC25A40 (C) in the K562 cells treated with 1mM BSO for 2d. ^*^, SLC25A40 band. A39 mRNA level does not change upon BSO treatment (B).

**Figure S2.**
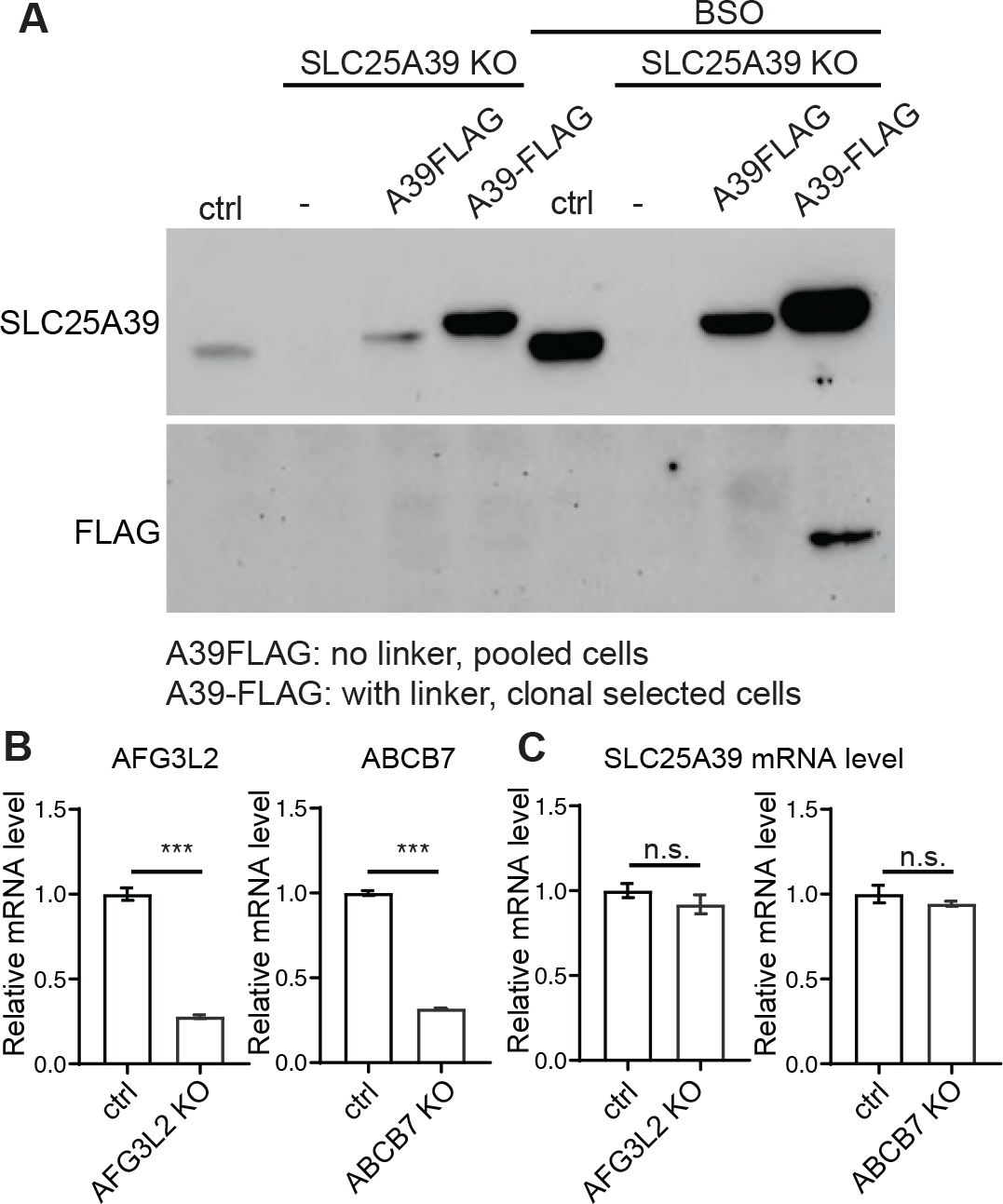
**(A)** Western blot of A39 and FLAG for the endogenous A39, ectopically expressed A39FLAG (without flexible linker), A39-FLAG (with flexible linker) in the A39 CRISPR KO K562 cells, with or without 1mM BSO treatment for 2d. (B) Validation of AFG3L2 and ABCB7 CRISPR KO K562 cells by reduced mRNA level through nonsense mediated decay. (C) A39 mRNA level is unchanged in the AFG3L2 and ABCB7 CRISPR KO cells.

**Figure S3.**
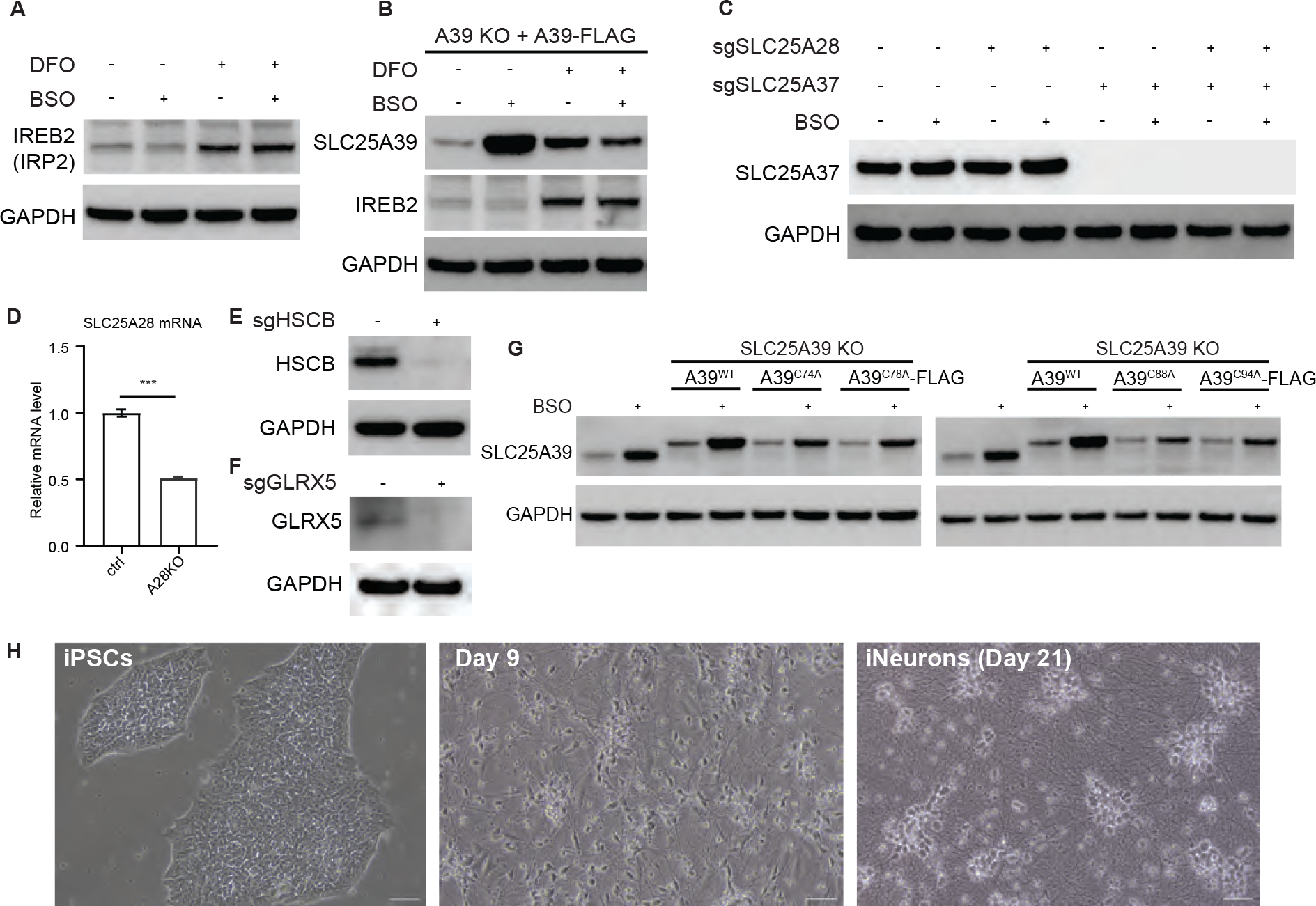
Validation of iron manipulation. (A) Western blot of IRP2 upon BSO (1mM, 2d) and DFO (20 μM, 2d) treatment. (B) Western blot of A39 and IRP2 from the A39 CRISPR KO K562 cells expressing A39-FLAG upon BSO (1mM, 2d) and DFO (20 μM, 2d) treatment. (C) Western blot of SLC25A37 from the control and SLC25A39 CRISPR KO K562 cells upon BSO treatment (1mM, 2d). (D) Validation of SLC25A28 CRISPR KO by reduced SLC25A28 mRNA level through nonsense mediated decay. Statistical significance was calculated using two-tailed *t* test. Significance level was indicated as ^***^ *p* < 0.001. Data are expressed as mean ± SD. (E-F) Western blot of HSCB (E) and GLRX5 (F) in the corresponding CRISPR KO K562 cells. (G) A39 western blot for the endogenous A39, ectopically expressed A39^WT^ and single cysteine mutants A39^C74A^, A39^C78A^, A39^C88A^, A39^C94A^ in the A39 CRISPR KO K562 cells upon BSO treatment (1mM, 2d). (H) Cell morphology of iPSCs, developing neurons (Day 9 post differentiation), and mature neurons (Day 21 post differentiation). Scale bar, 100 μm.

## References

1. Cunningham, C.N., and Rutter, J. (2020). 20,000 picometers under the OMM: diving into the vastness of mitochondrial metabolite transport. EMBO Rep 21, e50071. 10.15252/embr.202050071.

2. Kunji, E.R.S., King, M.S., Ruprecht, J.J., and Thangaratnarajah, C. (2020). The SLC25 Carrier Family: Important Transport Proteins in Mitochondrial Physiology and Pathology. Physiology (Bethesda) 35, 302–327. 10.1152/physiol.00009.2020.

3. Palmieri, F. (2013). The mitochondrial transporter family SLC25: identification, properties and physiopathology. Mol Aspects Med 34, 465–484. 10.1016/j.mam.2012.05.005.

4. Bar-Peled, L., and Kory, N. (2022). Principles and functions of metabolic compartmentalization. Nat Metab 4, 1232–1244. 10.1038/s42255-022-00645-2.

5. Balaban, R.S., Nemoto, S., and Finkel, T. (2005). Mitochondria, oxidants, and aging. Cell 120, 483–495. 10.1016/j.cell.2005.02.001.

6. Murphy, M.P. (2009). How mitochondria produce reactive oxygen species. Biochem J 417, 1–13. 10.1042/BJ20081386.

7. Finkel, T. (2012). Signal transduction by mitochondrial oxidants. J Biol Chem 287, 4434–4440. 10.1074/jbc.R111.271999.

8. D’Autreaux, B., and Toledano, M.B. (2007). ROS as signalling molecules: mechanisms that generate specificity in ROS homeostasis. Nat Rev Mol Cell Biol 8, 813–824. 10.1038/nrm2256.

9. Meister, A., and Anderson, M.E. (1983). Glutathione. Annu Rev Biochem 52, 711–760. 10.1146/annurev.bi.52.070183.003431.

10. Mari, M., Morales, A., Colell, A., Garcia-Ruiz, C., Kaplowitz, N., and Fernandez-Checa, J.C. (2013). Mitochondrial glutathione: features, regulation and role in disease. Biochim Biophys Acta 1830, 3317–3328. 10.1016/j.bbagen.2012.10.018.

11. Rouault, T.A., and Tong, W.H. (2008). Iron-sulfur cluster biogenesis and human disease. Trends Genet 24, 398–407. 10.1016/j.tig.2008.05.008.

12. Lill, R. (2009). Function and biogenesis of iron-sulphur proteins. Nature 460, 831–838. 10.1038/nature08301.

13. Wang, Y., Yen, F.S., Zhu, X.G., Timson, R.C., Weber, R., Xing, C., Liu, Y., Allwein, B., Luo, H., Yeh, H.W., et al. (2021). SLC25A39 is necessary for mitochondrial glutathione import in mammalian cells. Nature 599, 136–140. 10.1038/s41586-021-04025-w.

14. Shi, X., Reinstadler, B., Shah, H., To, T.L., Byrne, K., Summer, L., Calvo, S.E., Goldberger, O., Doench, J.G., Mootha, V.K., and Shen, H. (2022). Combinatorial GxGxE CRISPR screen identifies SLC25A39 in mitochondrial glutathione transport linking iron homeostasis to OXPHOS. Nat Commun 13, 2483. 10.1038/s41467-022-30126-9.

15. Shi, X., Reinstadler, B., Shah, H., To, T.-L., Byrne, K., Summer, L., Calvo, S.E., Goldberger, O., Doench, J.G., Mootha, V.K., and Shen, H. (2021). Combinatorial G x G x E CRISPR screening and functional analysis highlights SLC25A39 in mitochondrial GSH transport. bioRxiv, 2021.2009.2022.461361. 10.1101/2021.09.22.461361.

16. Byrne, K.L., Szeligowski, R.V., and Shen, H. (2023). Phylogenetic Analysis Guides Transporter Protein Deorphanization: A Case Study of the SLC25 Family of Mitochondrial Metabolite Transporters. Biomolecules 13. 10.3390/biom13091314.

17. Jumper, J., Evans, R., Pritzel, A., Green, T., Figurnov, M., Ronneberger, O., Tunyasuvunakool, K., Bates, R., Zidek, A., Potapenko, A., et al. (2021). Highly accurate protein structure prediction with AlphaFold. Nature 596, 583–589. 10.1038/s41586-021-03819-2.

18. Murphy, M.P. (2012). Mitochondrial thiols in antioxidant protection and redox signaling: distinct roles for glutathionylation and other thiol modifications. Antioxid Redox Signal 16, 476–495. 10.1089/ars.2011.4289.

19. Requejo, R., Hurd, T.R., Costa, N.J., and Murphy, M.P. (2010). Cysteine residues exposed on protein surfaces are the dominant intramitochondrial thiol and may protect against oxidative damage. FEBS J 277, 1465–1480. 10.1111/j.1742-4658.2010.07576.x.

20. Nilsson, B.L., Soellner, M.B., and Raines, R.T. (2005). Chemical synthesis of proteins. Annu Rev Biophys Biomol Struct 34, 91–118. 10.1146/annurev.biophys.34.040204.144700.

21. Cho, N.H., Cheveralls, K.C., Brunner, A.D., Kim, K., Michaelis, A.C., Raghavan, P., Kobayashi, H., Savy, L., Li, J.Y., Canaj, H., et al. (2022). OpenCell: Endogenous tagging for the cartography of human cellular organization. Science 375, eabi6983. 10.1126/science.abi6983.

22. Mathieson, T., Franken, H., Kosinski, J., Kurzawa, N., Zinn, N., Sweetman, G., Poeckel, D., Ratnu, V.S., Schramm, M., Becher, I., et al. (2018). Systematic analysis of protein turnover in primary cells. Nat Commun 9, 689. 10.1038/s41467-018-03106-1.

23. Zecha, J., Meng, C., Zolg, D.P., Samaras, P., Wilhelm, M., and Kuster, B. (2018). Peptide Level Turnover Measurements Enable the Study of Proteoform Dynamics. Mol Cell Proteomics 17, 974–992. 10.1074/mcp.RA118.000583.

24. Morgenstern, M., Peikert, C.D., Lubbert, P., Suppanz, I., Klemm, C., Alka, O., Steiert, C., Naumenko, N., Schendzielorz, A., Melchionda, L., et al. (2021). Quantitative highconfidence human mitochondrial proteome and its dynamics in cellular context. Cell Metab 33, 2464–2483 e2418. 10.1016/j.cmet.2021.11.001.

25. Chen, W.W., Freinkman, E., and Sabatini, D.M. (2017). Rapid immunopurification of mitochondria for metabolite profiling and absolute quantification of matrix metabolites. Nat Protoc 12, 2215–2231. 10.1038/nprot.2017.104.

26. Chen, W.W., Freinkman, E., Wang, T., Birsoy, K., and Sabatini, D.M. (2016). Absolute Quantification of Matrix Metabolites Reveals the Dynamics of Mitochondrial Metabolism. Cell 166, 1324–1337 e1311. 10.1016/j.cell.2016.07.040.

27. Horwich, A.L., Fenton, W.A., Chapman, E., and Farr, G.W. (2007). Two families of chaperonin: physiology and mechanism. Annu Rev Cell Dev Biol 23, 115–145. 10.1146/annurev.cellbio.23.090506.123555.

28. Arlt, H., Tauer, R., Feldmann, H., Neupert, W., and Langer, T. (1996). The YTA10-12 complex, an AAA protease with chaperone-like activity in the inner membrane of mitochondria. Cell 85, 875–885. 10.1016/s0092-8674(00)81271-4.

29. Di Bella, D., Lazzaro, F., Brusco, A., Plumari, M., Battaglia, G., Pastore, A., Finardi, A., Cagnoli, C., Tempia, F., Frontali, M., et al. (2010). Mutations in the mitochondrial protease gene AFG3L2 cause dominant hereditary ataxia SCA28. Nat Genet 42, 313–321. 10.1038/ng.544.

30. Pierson, T.M., Adams, D., Bonn, F., Martinelli, P., Cherukuri, P.F., Teer, J.K., Hansen, N.F., Cruz, P., Mullikin For The Nisc Comparative Sequencing Program, J.C., Blakesley, R.W., et al. (2011). Whole-exome sequencing identifies homozygous AFG3L2 mutations in a spastic ataxia-neuropathy syndrome linked to mitochondrial m-AAA proteases. PLoS Genet 7, e1002325. 10.1371/journal.pgen.1002325.

31. Srinivasan, V., Pierik, A.J., and Lill, R. (2014). Crystal structures of nucleotide-free and glutathione-bound mitochondrial ABC transporter Atm1. Science 343, 1137–1140. 10.1126/science.1246729.

32. Koppen, M., Metodiev, M.D., Casari, G., Rugarli, E.I., and Langer, T. (2007). Variable and tissue-specific subunit composition of mitochondrial m-AAA protease complexes linked to hereditary spastic paraplegia. Mol Cell Biol 27, 758–767. 10.1128/MCB.01470-06.

33. Atorino, L., Silvestri, L., Koppen, M., Cassina, L., Ballabio, A., Marconi, R., Langer, T., and Casari, G. (2003). Loss of m-AAA protease in mitochondria causes complex I deficiency and increased sensitivity to oxidative stress in hereditary spastic paraplegia. J Cell Biol 163, 777–787. 10.1083/jcb.200304112.

34. Patron, M., Tarasenko, D., Nolte, H., Kroczek, L., Ghosh, M., Ohba, Y., Lasarzewski, Y., Ahmadi, Z.A., Cabrera-Orefice, A., Eyiama, A., et al. (2022). Regulation of mitochondrial proteostasis by the proton gradient. EMBO J 41, e110476. 10.15252/embj.2021110476.

35. Luk, E., Carroll, M., Baker, M., and Culotta, V.C. (2003). Manganese activation of superoxide dismutase 2 in Saccharomyces cerevisiae requires MTM1, a member of the mitochondrial carrier family. Proc Natl Acad Sci U S A 100, 10353–10357. 10.1073/pnas.1632471100.

36. Nilsson, R., Schultz, I.J., Pierce, E.L., Soltis, K.A., Naranuntarat, A., Ward, D.M., Baughman, J.M., Paradkar, P.N., Kingsley, P.D., Culotta, V.C., et al. (2009). Discovery of genes essential for heme biosynthesis through large-scale gene expression analysis. Cell Metab 10, 119–130. 10.1016/j.cmet.2009.06.012.

37. Zhang, Y., Pak, C., Han, Y., Ahlenius, H., Zhang, Z., Chanda, S., Marro, S., Patzke, C., Acuna, C., Covy, J., et al. (2013). Rapid single-step induction of functional neurons from human pluripotent stem cells. Neuron 78, 785–798. 10.1016/j.neuron.2013.05.029.

38. Fernandopulle, M.S., Prestil, R., Grunseich, C., Wang, C., Gan, L., and Ward, M.E. (2018). Transcription Factor-Mediated Differentiation of Human iPSCs into Neurons. Curr Protoc Cell Biol 79, e51. 10.1002/cpcb.51.

39. Pantazis, C.B., Yang, A., Lara, E., McDonough, J.A., Blauwendraat, C., Peng, L., Oguro, H., Kanaujiya, J., Zou, J., Sebesta, D., et al. (2022). A reference human induced pluripotent stem cell line for large-scale collaborative studies. Cell Stem Cell 29, 1685–1702 e1622. 10.1016/j.stem.2022.11.004.

40. Sprenger, H.G., MacVicar, T., Bahat, A., Fiedler, K.U., Hermans, S., Ehrentraut, D., Ried, K., Milenkovic, D., Bonekamp, N., Larsson, N.G., et al. (2021). Cellular pyrimidine imbalance triggers mitochondrial DNA-dependent innate immunity. Nat Metab 3, 636–650. 10.1038/s42255-021-00385-9.

41. Quiros, P.M., Langer, T., and Lopez-Otin, C. (2015). New roles for mitochondrial proteases in health, ageing and disease. Nat Rev Mol Cell Biol 16, 345–359. 10.1038/nrm3984.

42. Liu, Y., Liu, S., Tomar, A., Yen, F.S., Unlu, G., Ropek, N., Weber, R.A., Wang, Y., Khan, A., Gad, M., et al. (2023). Autoregulatory control of mitochondrial glutathione homeostasis. Science, eadf4154. 10.1126/science.adf4154.

43. Magri, S., Fracasso, V., Plumari, M., Alfei, E., Ghezzi, D., Gellera, C., Rusmini, P., Poletti, A., Di Bella, D., Elia, A.E., et al. (2018). Concurrent AFG3L2 and SPG7 mutations associated with syndromic parkinsonism and optic atrophy with aberrant OPA1 processing and mitochondrial network fragmentation. Hum Mutat 39, 2060–2071. 10.1002/humu.23658.

